# Proto-ribosome: a theoretical approach based on RNA relics

**DOI:** 10.1101/182055

**Authors:** Jacques Demongeot

## Abstract

We describe in this paper, based on already published articles, a contribution to the theory postulating the existence of a proto-ribosome, which could have appeared early at the origin of life and we discuss the interest of this notion in an evolutionary perspective, taking into account the existence of possible RNA relics of this proto-ribosome.

---

Ten years after the first observations of a ribosome sixty years ago by G. E. Palade and P. Siekevit (*1*), theoreticians elaborated the first models on the ribosomal translation mechanism using tools coming from statistical mechanics and physical chemistry (*2*-*4*). Simultaneously, mathematicians like S. Ulam and J. Conway simulated large automata networks and remarked that with simple rules, they obtained complex numerical behaviours similar to those observed in biology, like motion and division (*5, 6*). Then, biologists defined the notions of genetic code ancestor and proto-ribosome, regularly used after by numerous authors (*7*-*27*) until some recent papers on the evolutionary biology (*28, 29*). We will give in the following the essential of three papers treating this topic in the wider framework of genetic code and biological boundaries (*30*-*32*).

## The genetic code as optimizing a variational principle

The genetic code consists in dispatching 64 triplets made of 3 letters representing purine bases – A for Adenine and G for Guanine - and pyrimidine ones – U for Uracil and C for Cytosine – into 21 synonymy classes. Each class can contain between 1 and 6 triplets, the 20 classes corresponding to the 20 amino acids (except for one class containing only 1 triplet, which corresponds either to the amino acid Methionine or, if this triplet initiates a sequence of messenger RNA (mRNA), to a “start” punctuation symbol), plus one class corresponding to the “end” punctuation symbol finishing the mRNA sequences. It has been proved that stereo-chemical bonds can favor a non-permanent reversible link between amino acids and triplets of their synonymy class *(33-35),* despite some criticisms *(36).*

An archetypal circular RNA structure called AL (for Archetypal Loop) exists obeying following opposite constraints, *i.e.*, satisfying a **min-max** problem:

- to be as short as possible,
- to present in its sequence at least one triplet corresponding to each amino acid, in order to serve as “matrimonial agency” favoring the vicinity of any couple of amino acids present in the original soup, close to the circular RNA AL, and able to form a strong peptide bond (*i.e.*, the covalent chemical bond formed by two amino acids, when the carboxyl group of one reacts with the amino group of the other) between them, in order to initiate the peptide building, such as an ancestral ribosome. For satisfying this constraint, the circular RNA AL must contain at least 20 triplets.

By solving the **min-max** problem above, we find the structure of Fig. 1 (*11*, *13*, *18*, *21*). This structure can be obtained in a circular or hairpin form and could be considered as the ancestor of the present ribosomes.

**Fig. 1.**
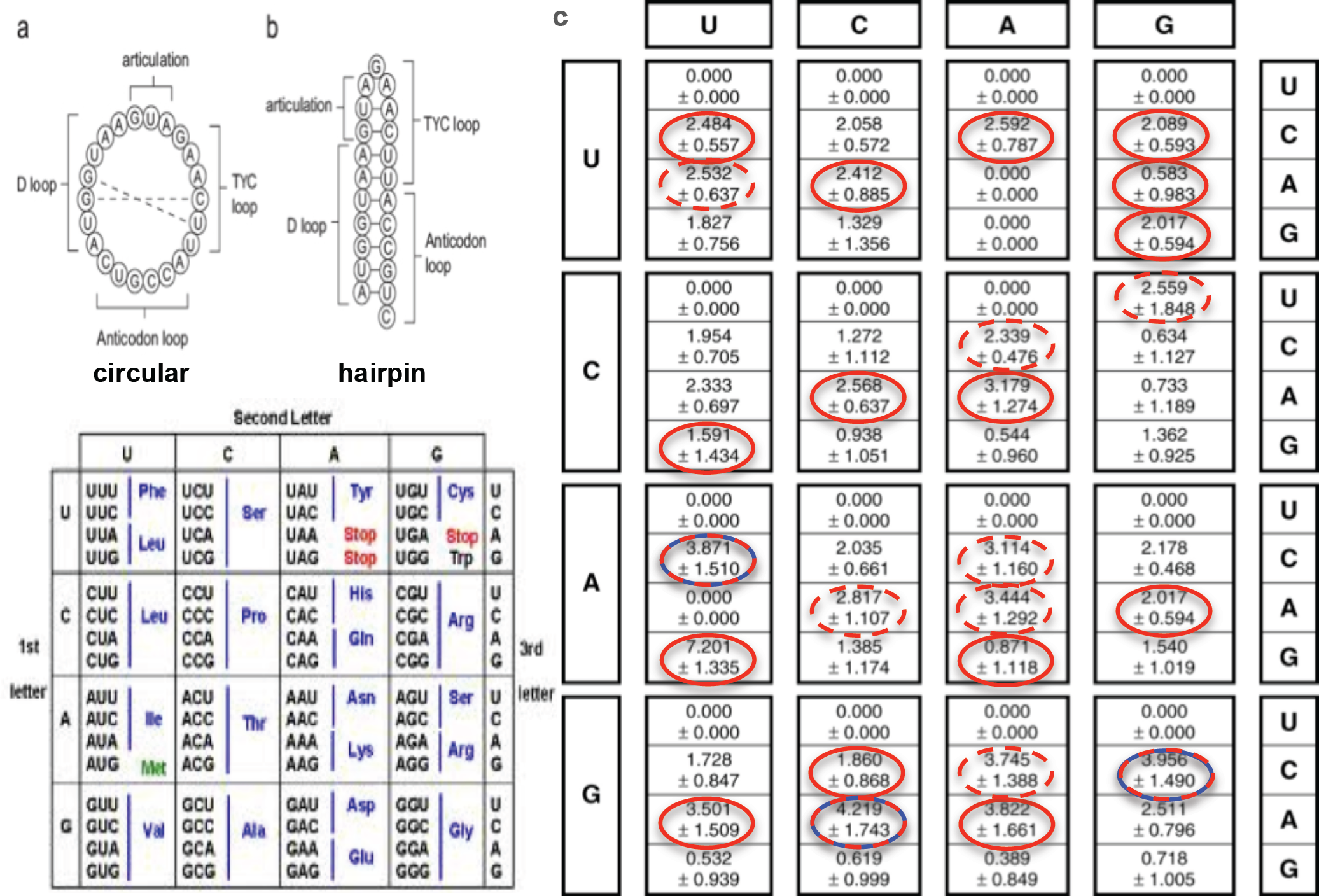
The AL structure is given in (a) circular and (b) hairpin forms. From (*37*), we have extracted the triplets which are present (c) in the sequence of the structure AL (3’-5’) and have the maximum frequency in a set of ^t^RNAs from 29 selected organisms with the corresponding standard deviations (red ellipses), or in the reverse sequence C’ (5’-3’) (red dotted ellipses), or in the anti-strand sequence C’’ (blue-red ellipses). The 22 AL triplets (with overlap) correspond to all the synonymy classes (d).

It is possible to show (*30*-*32*) that the codon repartition of the genetic code satisfies a variational principle equalling the normalized mean error **M** due to mutation between 2 synonymy classes of frequency f and (1-f) respectively and the normalized information **I** of the Bernoulli distribution B(f,1-f) defined by:

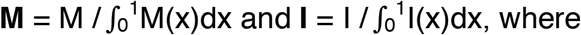

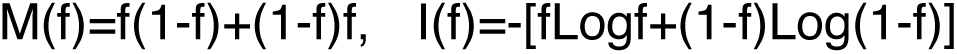, 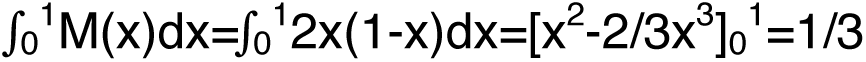, 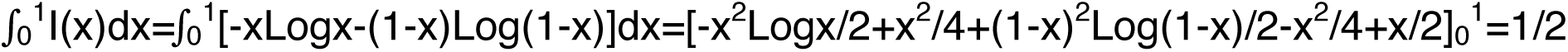.

The normalized functions **M** and **I**, whose integral equal 1, verify (Fig. 2 bottom left):

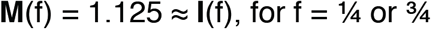

Hence, we obtain the repartition R of triplets inside the genetic code (Fig. 1d) after 10 successive partitions into 2 subclasses respecting the optimal frequency f=1/4 (Fig. 2).

**Fig. 2.**
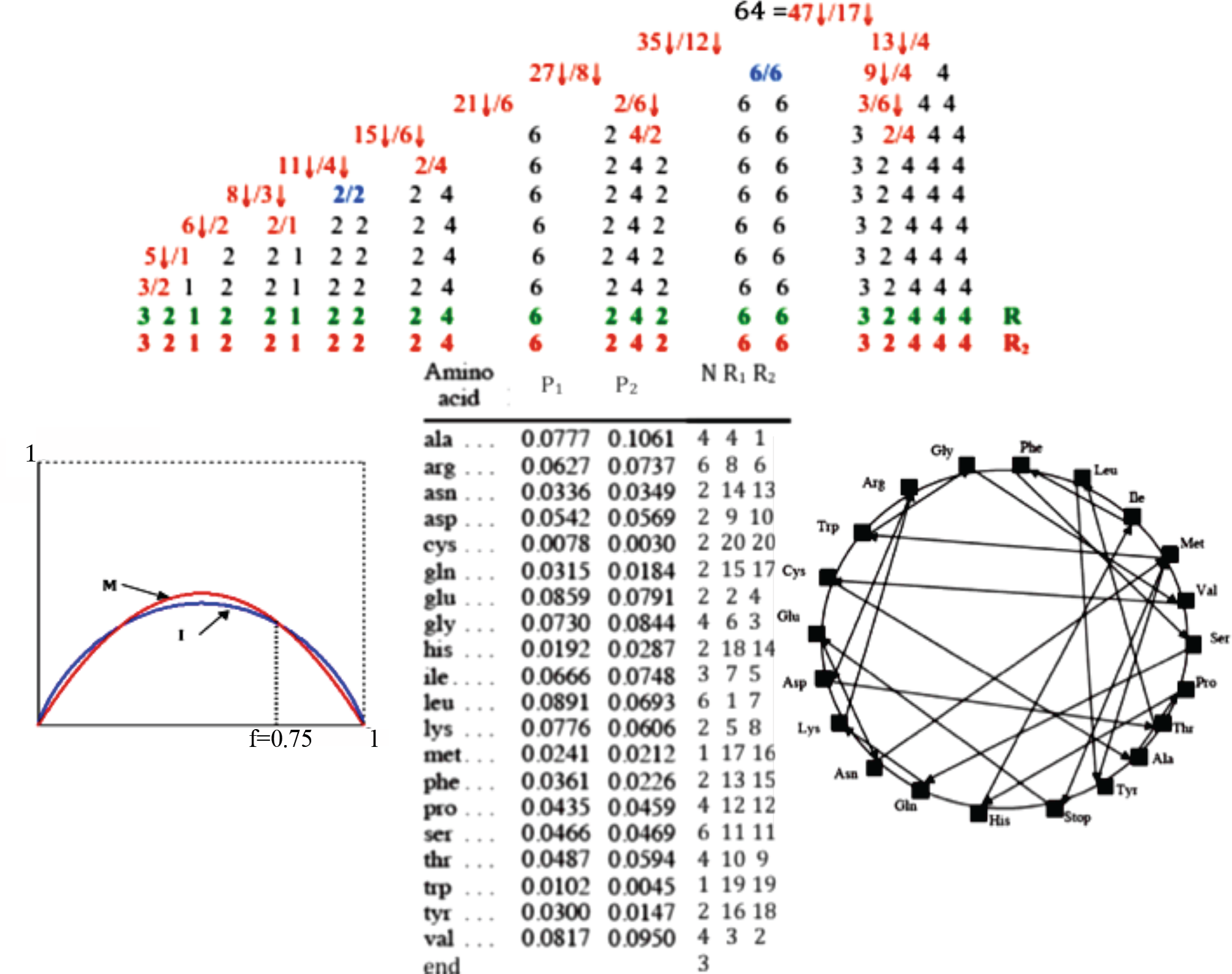
Top: Successive repartition of the 64 codons following the optimal rule of affectation: when the ratio between sizes of the smaller and larger new classes is about 3 it is indicated in red. This optimal repartition R of amino acids respects at each division of triplet classes the min-max rule equalling the information ***I***, and the mutational behaviour ***M***, *i.e.*, about the proportions 3/4 – 1/4. Bottom left: graph of the functions M and I with indication of the optimal f. Bottom middle: ranking of the amino acids depending on their frequency in proteins of 8 species (P_1_) and in their Last Universal Ancestor (LUA) (P_2_); N denotes the size of the amino acid class, R_1_ its ranking based on P_1_ and R_2_ on P_2_. R_2_ is the same as the ranking R obtained from the optimal repartition (*38*). Bottom right: Hamiltonian path solving the variational problem on a graph whose vertices represent the 21 synonymy classes, counting twice Met.

The main property of the circular RNA sequence AL is to include the most frequent triplets (Fig. 1) in ^t^RNAs of 29 selected organisms (*39*), allowing revisiting amino acid ranking from their appearance order in various experiments, which are all coherent, giving the same ranking for the first 6 amino acids occurring in a neo-synthesis of Miller’s type (Table 1).

**Table 1.**
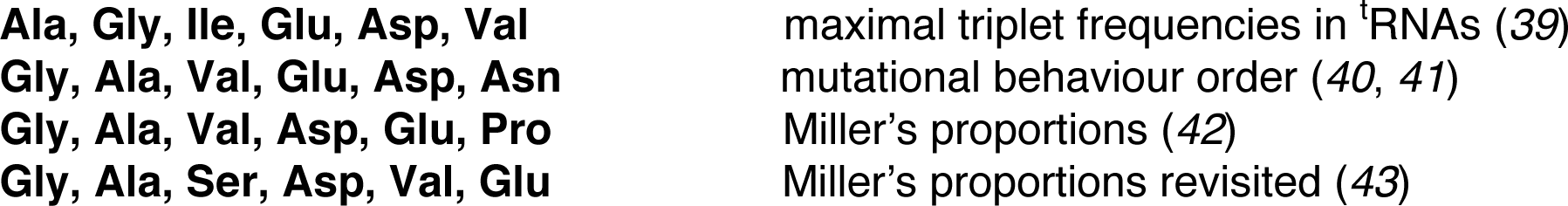
Amino acids ranking (except for Met/Start) in various sets of nucleic acids from different organisms and experiments of RNA neo-synthesis

**Fig. 3.**
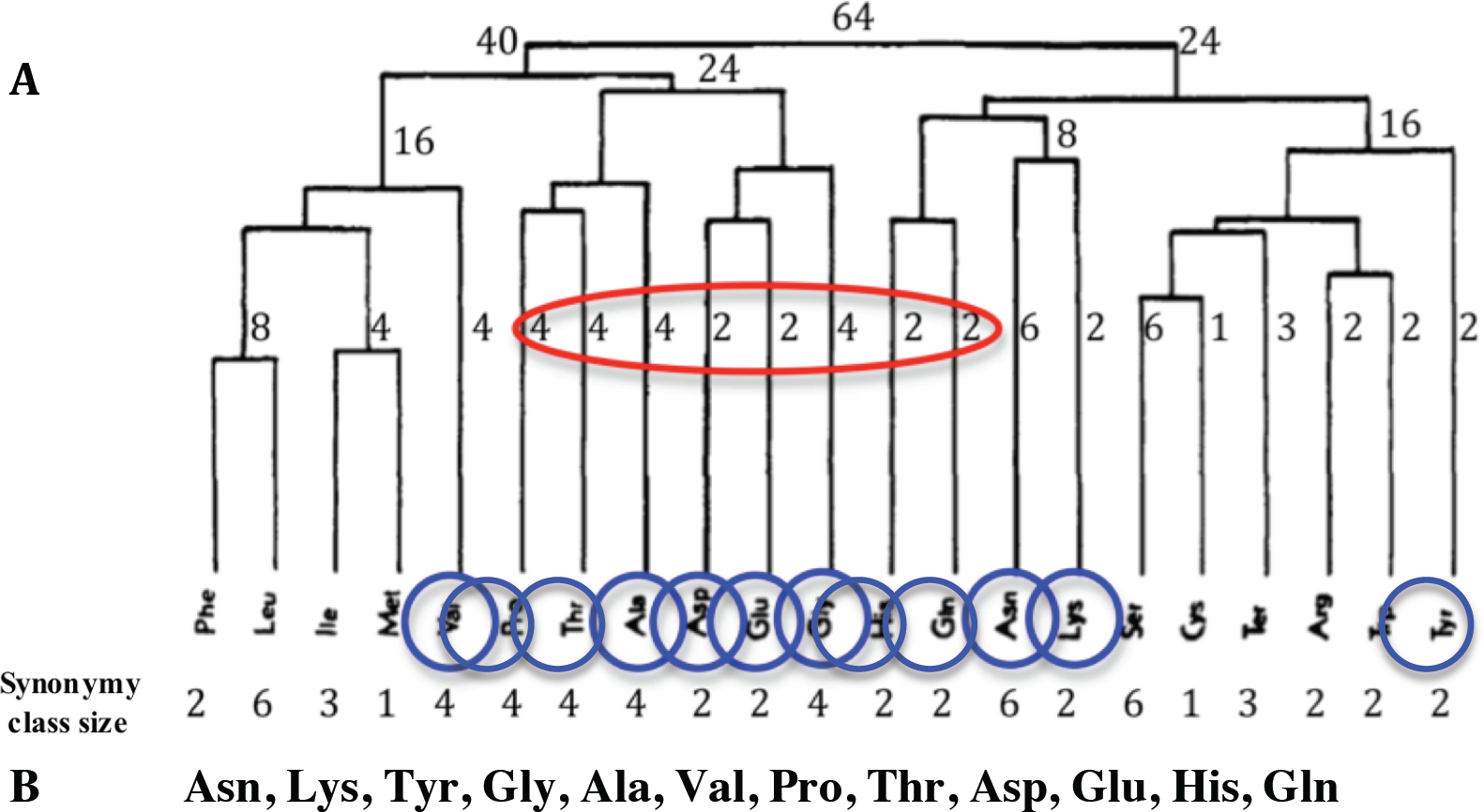
A: The amino acids appearing progressively in a clustering based on their mutational behaviour, with indication of the synonymy class sizes. B: Order of the first amino acids appearing at the top of the dendrogram, whose start clusters (in red) are similar to the beginning of the optimal rule (Fig. 2 top).

The amino acids appear in Fig. 2 in a hierarchical clustering process based on their mutational behaviour proposed in (*40*), and the progressive concatenation of clusters follows approximately the optimal repartition of amino acids respecting, at each triplet class division, a **min-max** rule consisting of maximizing the entropy and minimizing the mutational behaviour of amino acids, depending on their synonymy class size. If we rank the amino acids (Table 1 and Fig. 4) by using either their relative concentration in Miller’s experiment (*42*) thought to reproduce the conditions of the prebiotic earth (*44*), or their codons’ thermo-stability (*45*), or their concentrations in revisited Miller’s data (*43*).

**Fig. 4.**
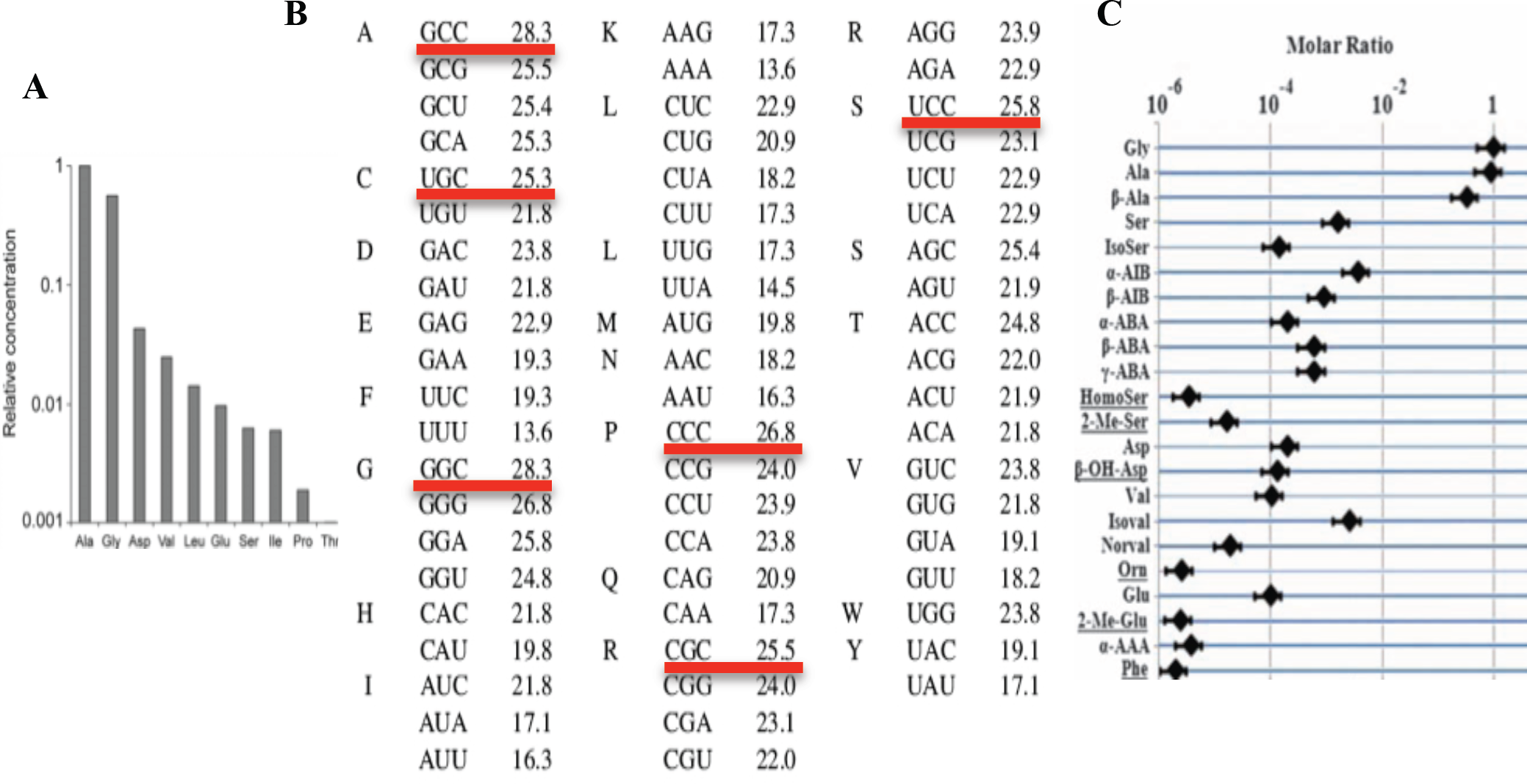
A. Amino acids ranking by using their relative concentration in Miller’s experiment (*44*) B. Codons thermo-stability (kcal/M) (*45*) C. Amino acids concentrations in revisited Miller’s data (*43*).

## Biological boundaries

A cell can be defined anatomically by its membranes and functionally by the concentration or gradient boundaries of the metabolites involved in its main survival functions (Fig. 5). The gradient boundary of a metabolite is the line on which the mean Gaussian curvature C on the surface defined by its concentration c vanishes:

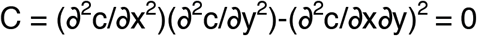

The k-concentration boundary is the contour line of the concentration surface on which the concentration c=k. The advantage of the gradient boundary over the concentration boundary comes from the fact that if the concentration surface is an isotropic Gaussian bell-shaped surface (for example in the case of isotropic diffusion of a metabolite from its source of production), then the gradient boundary is the zero-Laplacian curve, where diffusion vanishes: on such a boundary the metabolite can self-assembles or assembles with other membrane constituents if they are also slightly diffusing and if their co-existence time at the same place is sufficiently long for allowing this assemblage. We give in the following an example of use of the gradient boundary for the "proto-cells" at the origin of life.

## Proto-cell boundary

"A boundary is needed to separate life from non-life" (*46*). At the beginning of life, without any enzymes, RNA strands bound to longer template strands of RNA could grow in single nucleotide steps according to the base pairing rules of Watson and Crick. These duplication steps could take place on the timescale of hours (*47*) giving support to the theory that RNA spontaneously replicated during prebiotic evolution. Let us suppose now that the initial RNA template strands polymerized by chance in the prebiotic RNA world were circular with nucleotide sequences similar to those of a the Archetypal Loop (AL). This RNA ring offers weak interaction sites (electrostatic at short distance and van der Waals at mean distance) for any hydrophilic or hydrophobic amino acid present in the prebiotic medium. "The most obvious function of RNA today is to serve as a structural element that assists in the formation of strong peptide bonds between amino acids in the synthesis of proteins. The first RNAs may have served the same purpose, but without any preference for specific amino acids. Many further steps in evolution would be needed to "invent" the elaborate mechanisms for replication and specific protein synthesis that we observe in life today" (*46*). Such primitive RNA-dependent protein-building machinery (a "matrimonial agency" for amino acids) could have been more efficient than thioesters of amino acids which spontaneously form peptides in aqueous solutions (*48*) and would have played the same role as a ribosome, the machine that presently makes the proteins inside cells. The high similarity between AL and the sequence made of the concatenation of the ^t^RNA loops (D-, anticodon- and Ty-loops) reinforces this hypothesis (cf. *13*, *18*, *21* and AL supplementary material one).

The boundary of the first functional "proto-cell" able to build peptides can be defined as a peptide gradient boundary centred on the "proto-ribosome" AL, resulting from an a-specific confinement of amino acids around AL favouring the occurrence of peptide bounds. This "proto-ribosome" into a "proto-membrane" constituted the "proto-cell" with a circular organization satisfying a variational principle: peptide synthesis favoured by AL was necessary to build and repair the membrane of the proto-cell, made of hydrophobic peptides ensuring the integrity of the RNA AL "proto-ribosome", hence protected against denaturation, like in a principle of parsimony "à la Maupertuis" (*49*).

**Fig. 5.**
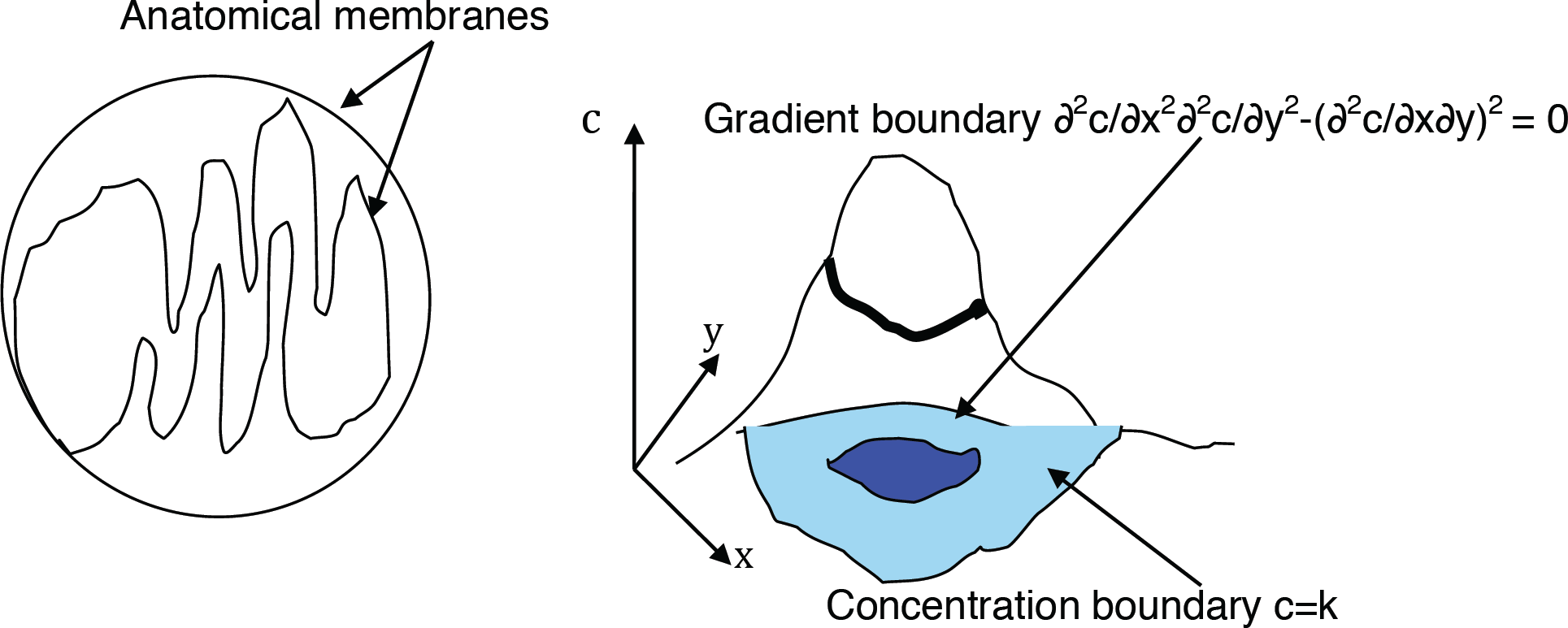
Anatomical (left) and functional boundaries (right).

The scenario presented above is similar to the behaviour of the auto-poietic automaton studied in (*50*-*52*), which is in exponential growth if the proto-membrane allows the entry of nucleic bases permitting AL as well as anti-sense AL replication, anti-sense AL being very similar to AL because of its internal symmetry (Fig. 6). The ring AL (respectively the ring anti-sense AL) selects and confines L-amino acids (respectively D-amino acids), and builds either hydrophobic or hydrophilic peptides (*53*) depending on the pole used for the confinement, acting as a proto-ribosome. In Fig. 6 top left, we indicate the AL hydrophobic (respectively hydrophilic) pole, where codons bind mainly hydrophobic (respectively hydrophilic) amino acids, the excess of the polar hydrophobicity (respectively hydrophilicity) being equal to 13.4 (resp. -21.3) (*54*).

According to the wobble hypothesis by Crick, the first two bases of the codons are essential for their assignation to a given amino acid and recognition by their ^t^RNA anti-codon (*55*): like in (*56*, *57*), we observed that the second base of the codons permits to associate them with hydrophobic amino acids, if this second base is Uracil U or Cytosine C and the first base is Guanine G, and with hydrophilic amino acids, if this second base is Adenine A or Cytosine C and the first base is either Adenine A or Uracil U (*54*). Cytosine C is difficult to obtain at the origin of life (*58*-*60*), but only 5 amino acids were not confined by AL without C, these missing amino acids being Alanine, Proline, Threonine, Glutamine and Histidine, respectively the 3^rd^, 7^th^, 10^th^, 12^th^ and 17^th^ amino acids in the decreasing order of use in human proteins (*61*), showing that the most frequent amino acids can be associated to primitive RNA without the help of Cytosine.

Growth is stopped due to a lack of nucleic or protein "stones", *i.e.*, because of the exhaustion of the basic atoms C, H, O, N, whose pool gives amino acids and nucleotides during the exponential growth, like in the Miller's experiment (*42*). If we denote by R (resp. A, B, M and P) the AL or anti-sense AL **R**ing (resp. **A**mino acids, nucleotide **B**ases, hydrophobic peptide **M**embranes and C,H,O,N-**P**ool) concentration, we have:

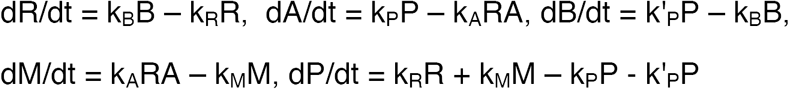

The differential system above presents an initial exponential growth behaviour and tends to a stationary state, if P is constant. If we add a diffusion term for the different metabolites above, this dynamics leads to the spatial segregation of R, A, B, M, P into structures like a “proto-nucleus” (R), “proto-cytoplasm” (A + B), “proto-membrane” (M) and building blocks (P). AL serves as a template for the formation of hydrophilic **E**nzymatic peptides E able to catalyse the AL or anti-sense AL replication (respectively degradation) with a reaction constant k_B_ (respectively k_R_) (*62*).

During its exponential growth, the boundary of the automaton is chosen as the gradient boundary of peptides polymerized from the 15 amino acids, that is all of them minus Alanine, Proline, Threonine, Glutamine and Histidine.

**Fig. 6.**
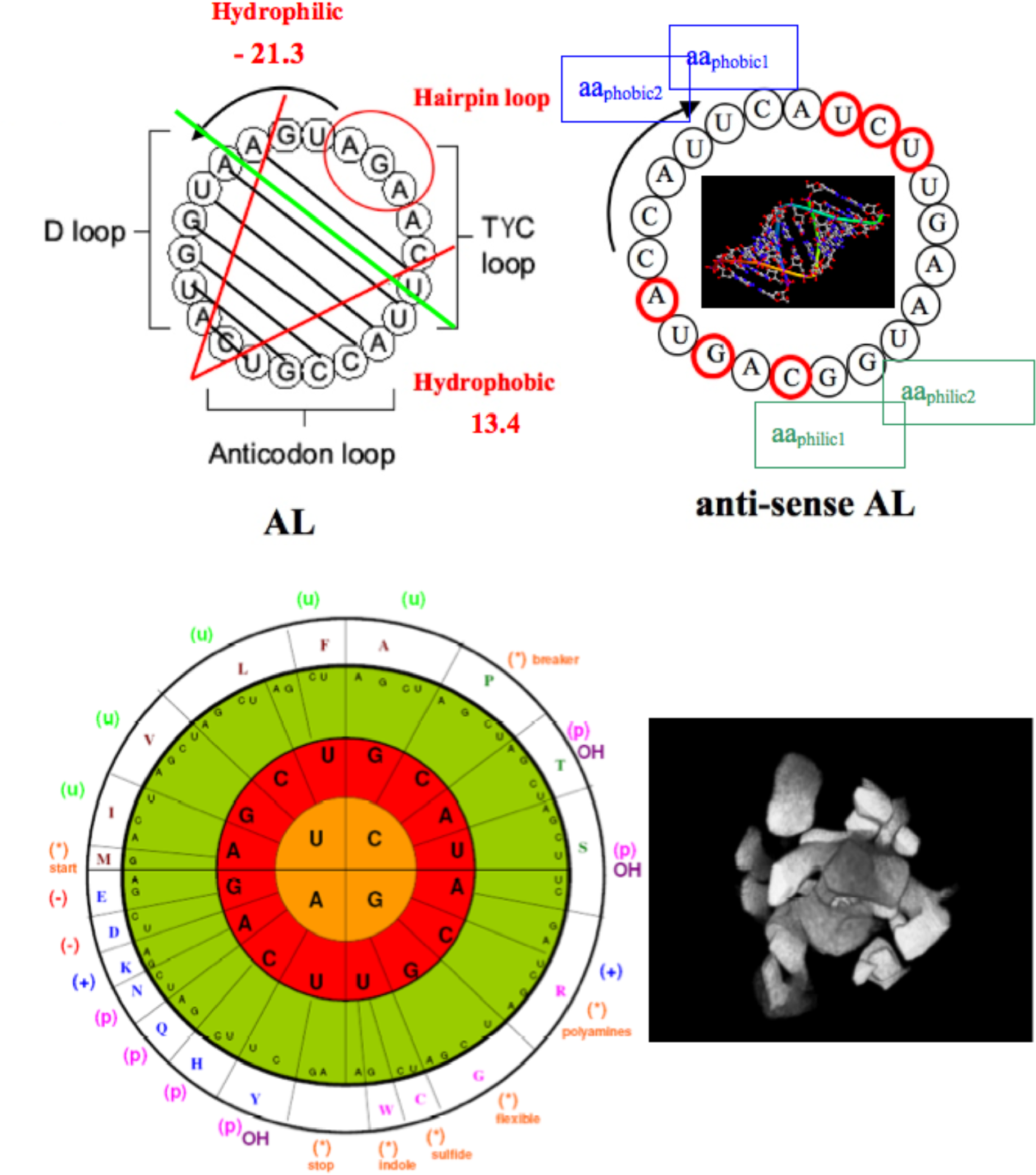
Top left: The RNA ring AL (with hydrophobic and hydrophilic poles). Top right: anti-sense AL. Bottom left: Coherence with the genetic code representation giving the hydrophobic (u) and hydrophilic (p) codons (*56*) showing that the AL order of bases roughly preserves the order of the first codon bases in the genetic code (red circle). Bottom right: Gradient boundaries of the M molecules showing “proto-membranes” of “proto-cell” structures (*63*).

**Fig. 7.**
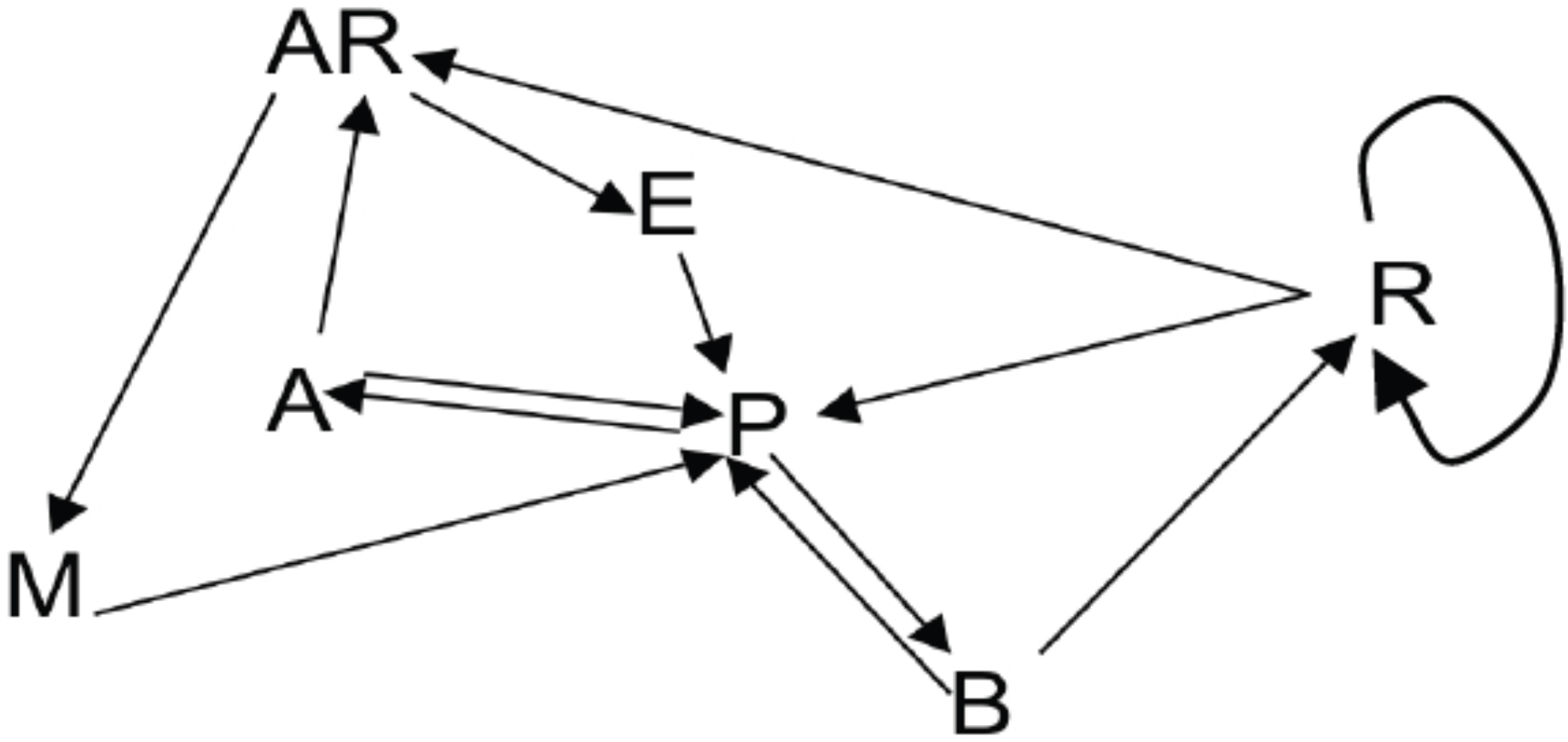
Primitive regulatory network of a “proto-cell” with only activatory arrows, showing different elements in interaction like a RNA proto-ribosome (R), proto-cytoplasmic constituents (A, Amino acids and B, nucleic Bases), a proto-Membrane (M), functional molecules such as Active RNA (AR) and Enzymes (E) and building blocks Products (P).

If we introduce diffusion (whose viscosity depends on the membrane concentration M) in the purely reaction differential system above, we get the new system below, having the graph of Fig. 7 as Jacobian graph, and whose discrete analogue has been simulated in (63) showing a progressive segmentation of the space by M molecules gradient:

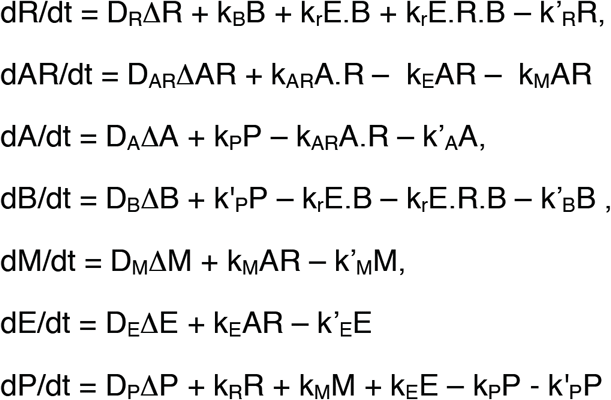

In case of isotropic diffusion, the zero-Laplacian (or zero-curvature or maximal gradient) line of the concentration surface of M molecules constitutes a gradient boundary and can become an anatomical frontier, if M auto-assembles with lipids. This line corresponds to the region where the mean Gaussian curvature of the concentration surface, defined by ∂^2^M/∂x^2^∂^2^M/∂y^2^ - (∂^2^M/∂x∂y)^2^, vanishes.

**Fig. 8.**
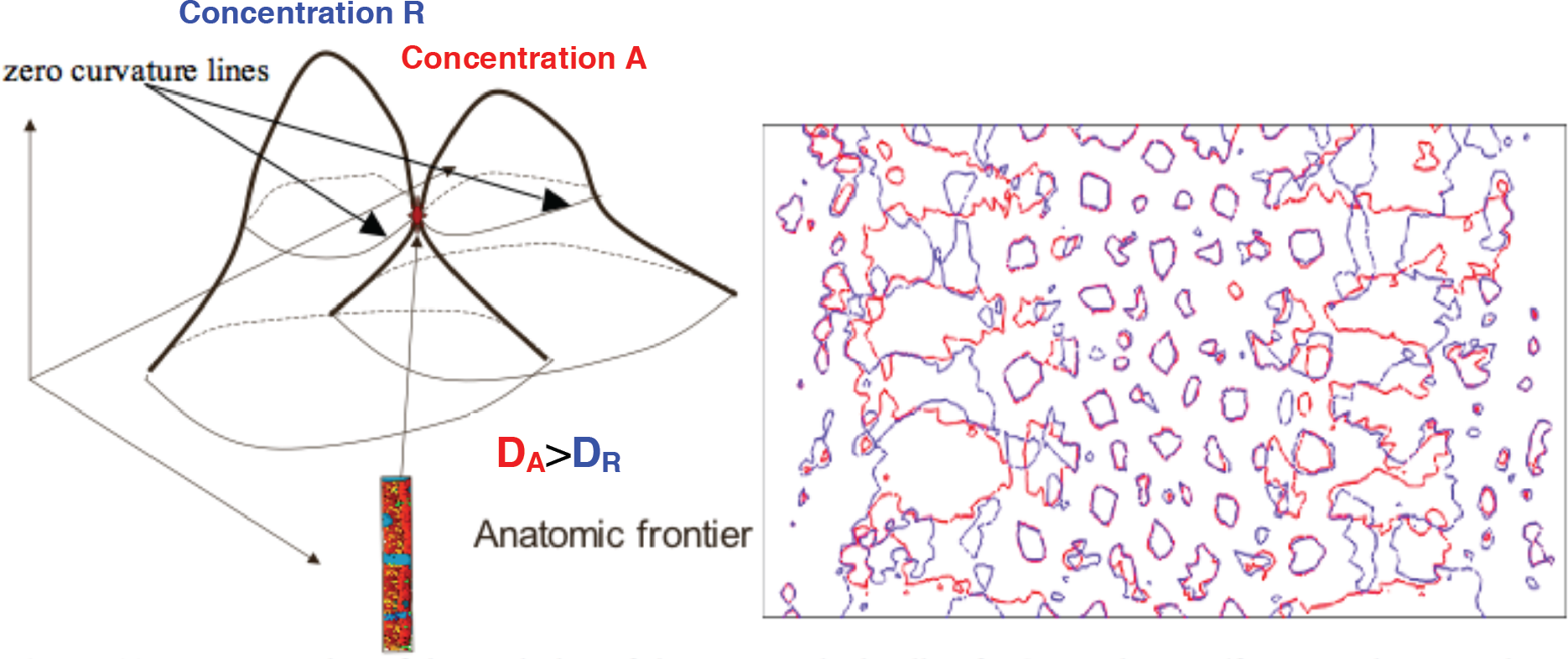
**Left:** Representation of the evolution of the zero-Laplacian line for 2 constituents R and A. Right: asymptotic co-existence of R (in blue) and A (in red) on the zero curvature (and diffusion) lines, allowing locally the assemblage of a “proto-membrane” (*64*).

If the biosynthesis of M results from the meeting between the two wave fronts made respectively of R and A molecules, let us consider the intersection of the zero-Laplacian lines of the corresponding concentration surfaces (Fig. 8): if the diffusion of R (respectively A) is isotropic from its initial synthesis at an initial P pool source, the R (respectively A) wave front progresses with the same velocity in all directions and the gradient boundary is the geometric locus where the diffusion in these directions vanishes. Therefore, at the intersection of the gradient boundaries, the co-existence time of R and A is maximum, favouring the amino acids / nucleic acids transient stereo-interaction, allowing the local peptide polymerization. Fig. 8 shows the possibility of such an intersection on only one tangency point or two intersection points (left) and on the whole gradient boundaries asymptotically confounded (right) for a convenient value of the ratio between the diffusion coefficients D_R_/D_A_ (*65*).

## Cell boundary

A hydrophobic proto-membrane plays the same role as a cell boundary made of amphiphilic lipid bilayers with membrane proteins. Peptides M interact with primitive amphiphilic components (*66*-*68*) or interact only with themselves like in co-cross-linked peptide artificial membranes (*69*).

ATP is present near the “proto-cells” because in an environment "poor in water but rich in ADP and inorganic phosphate, the formation of ATP is spontaneous, and ATP has a lower "energy" than ADP and inorganic phosphate" (*70*). The “protomembranes” could actively favour the penetration of ATP inside proto-cells if among the M peptides there exists membrane protenoids like the translocators present in micro-organisms, whose gene sequence matches with AL: the gene of the 2-oxoglutarate translocator of *Chlamydia muridarum* contains for example 6 AL heptamers for 1416 bpi, the expected number X being equal to 1409x22/2^14^ = 1.89±2.27*. The symbol * indicates the 95% confidence interval and the probability p of the observed number can be calculated from the normal approximation N(µ,s) of 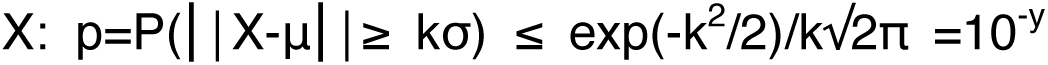, where y≈0.4+Log_10_k+0.217k^2^. Here k≈3, y≈3 and p≈10^-^3. Among the 6 AL heptamers, GAATGGT, which is part of the AL hemi-hairpin ATGAATGGTA, is observed twice, but the expected number is 0.086±0.48*, which corresponds to p=10^-^10. In *Encephalitozoon romaleae*, the observed number of AL heptamers in the ATP/ADP translocase is 9, with 2±2.5* expected, which correspond to p=10^-^5, and GAATGGT is also observed twice. Translocators (tlc) are considered as ancestors of the present translocases (*71*, *72*) and they could have been stereo-chemically synthesized from the AL template acting as a “proto-ribosome” (cf. AL supplementary material two).

## AL and nucleo-peptidic mechanisms

Different intracellular mechanisms involving RNA, DNA and proteins could conserve as relics sub-sequences of AL, in particular from its hemi-hairpin ATGAATGGTA.

## - Zinc finger mechanism

A zinc finger is a small protein serving as interaction module binding RNA, DNA and proteins, thanks to one or more zinc ions ensuring the stabilization of the molecular fold. Numerous zinc finger proteins in many species contain AL pentamers. For example, the gene of the mRNA of the zinc finger protein 84-like of *Jaculus jaculus* contains 180 AL pentamers for 2687 bpi (p=10^-^53) and 32 AL heptamers ATGAATG, for 0.16±0.7* expected, corresponding to p=10^-^1390 (cf. AL supplementary material three).

## - Polymerase mechanism

DNA (resp. RNA) polymerases are enzymes which synthesize DNA (resp. RNA) molecules from the nucleotides, ensuring the nucleic acids replication. They appeared very early in the evolution. The *Jaculus jaculus* polymerase alpha 1 contains for example 160 AL pentamers for 94±16* expected (p=10^-^11) and 5 AL heptamers AAGATGA for 0.25±0.85* expected, corresponding to p=10^-^20 (cf. AL supplementary material four).

## - Defence mechanism

The CRISPR-CAS system provides bacteria like *Streptococcus agalactiae* with adaptive immunity and we can notice that the AL pentamers ATGGT and ATTCA, and AL hexamers AATGGT and TCAAGAT (corresponding respectively to the D-loop TΨ-loop of many tRNAs) are often observed at many levels of this system (CASproteins, Casposon TIR and CRISP repeats) (cf. (*81*) and AL supplementary research five). For example, the typical repeat sequences for CRISPR1 and CRISPR3 (*73*) contain AL heptamers from ^t^RNA loops:

GTTTTTGTACTCTCAAGATTTAAGTAACTGTACAAC(CRISPR1)

GTTTTAGAGCTGTGTTGTTTCGAATGGTTCCAAAAC (CRISPR3)

as well as the sequences of TIR and CRISPR compared in (*74*), a consensus sequence from central part of the murine RSS VκL8, Jß2.6 and Jß2.2 (*75*-*77*), and the human RSS spacer common for Vh, V328h2 and V328 (*78*-*80*):

3’-ATACATCCC(C)TCTTAAGTTCCCTT-5’ (TIR)

3’-TTCCATCCC -TCTTAAGTTCGATT-5’ (CRISPR)

5’-ATGGTACTG - CCATTCAAGATGA-3’ (AL)

5’-GTGATACAG - CCCTTAACAAAAA-3’ (murine consensus RSS)

5’-ATTCAACATGAA-3’ (human RSS spacer)

The probability p=2 10^-^9 for 19 matches (with an insertion) between TIR and CRISPR using the binomial distribution B(1/4,22), p=8 10^-^6 for 15 anti-matches between AL and CRISPR plus 1 quasi-anti-match G-T using the distribution B(1/4,21)xB(3/8,1), p=7 10^-^4 for 13 matches between AL and consensus RSS using the binomial distribution B(1/4,22), p=2 10^-^6 for 11 matches between AL and RSS spacer using the binomial distribution B(1/4,12).

## Conclusion

To conclude, a small circular RNA, called AL, which presents the following features:

- its sub-sequences are observed as relics in many parts of the present genomes, namely in viral and non coding genomes (*24*),

- AL relics are often present in ^t^RNA loops (cf. (*18*, *25*) and AL supplementary material one),

- AL heptamer constitute the major part of the exon/intron boundary (cf. (*82*) and AL supplementary material five).

Hence, AL could have played the role of an ancient proto-ribosome: this claim is central in the stereochemical hypothesis of the genetic code formulated by A. Katchalsky in 1973 (*8*): the existence of catalytic RNAs in clays such as the “montmorillonite” may have facilitated the appearance of small peptides, involved secondarily (as now the protein replicase) in the replication of the RNA molecules, hence constituting a virtuous loop at the origin of life. The existence of a simple RNA structure capable to survive as a stable hairpin or to be functional in a ring form has been postulated early after Katchalsky’s hypothesis (*23*, *29*, *33*), and numerous experimental works (*34, 35*) reinforce this stereo-chemical hypothesis, despite criticisms (*36*), showing that the subject is still open, experimentally and theoretically.

## Acknowledgments

We are very indebted to J. Besson, A. Henrion-Caude, A. Moreira and P. Tracqui for many discussions about the molecular structures potentially involved at the origin of life.

The data reported in the paper are archived on the sites GtRNAdb, 5SRNAdb, GenBank, Protein Data Bank, Bacterial 16S rRNA Bioproject, Circbase, NCBI Blast and Kinefold or available in supplementary materials. The sites used are referenced as in the following:

